# Seq2MAIT: A Novel Deep Learning Framework for Identifying Mucosal Associated Invariant T (MAIT) Cells

**DOI:** 10.1101/2024.03.12.584395

**Authors:** Hesham ElAbd, Rachel Byron, Steven Woodhouse, Brittney Robinett, Joe Sulc, Andre Franke, Mitchell Pesesky, Wenyu Zhou, Haiyin Chen-Harris, Bryan Howie, Ruth Taniguchi, Harlan Robins

## Abstract

Mucosal-associated invariant T (MAIT) cells are a group of unconventional T cells that mainly recognize bacterial vitamin B metabolites presented on MHC-related protein 1 (MR1). MAIT cells have been shown to play an important role in controlling bacterial infection and in responding to viral infections. Furthermore, MAIT cells have been implicated in different chronic inflammatory diseases such as inflammatory bowel disease and multiple sclerosis. Despite their involvement in different physiological and pathological processes, a deeper understanding of MAIT cells is still lacking. Arguably, this can be attributed to the difficulty of quantifying and measuring MAIT cells in different biological samples which is commonly done using flow cytometry-based methods and single-cell-based RNA sequencing techniques. These methods mostly require fresh samples which are difficult to obtain, especially from tissues, have low to medium throughput, and are costly and labor-intensive. To address these limitations, we developed sequence-to-MAIT (*Seq2MAIT*) which is a transformer-based deep neural network capable of identifying MAIT cells in bulk TCR-sequencing datasets, enabling the quantification of MAIT cells from any biological materials where human DNA is available. Benchmarking *Seq2MAIT* across different test datasets showed an average area-under-the-receiver-operator-curve (AU[ROC]) >0.80. In conclusion, *Seq2MAIT* is a novel, economical, and scalable method for identifying and quantifying MAIT cells in virtually any biological sample.

## Introduction

The immune system in jawed vertebrates has been broadly categorized into innate and adaptive components. Innate immunity utilizes pattern recognition receptors such as toll-like receptors (TLR) and NOD-like receptors (NLR) to identify conserved markers of microbial infection such as peptidoglycans and lipopolysaccharides. Conversely, adaptive immunity uses V(D)J recombination, a somatic recombination mechanism that takes place on T and B cells, to generate a large repertoire of antigen-specific T and B cell receptors (TCR/BCR). A growing body of research has identified different groups of innate-like T and B cells termed innate-like or unconventional T and B cells, respectively^1–6^. Unconventional T cells, as opposed to conventional T cells, do not recognize peptides presented on polymorphic major histocompatibility complexes (MHC) and instead recognize lipids, glycolipids, metabolites, and peptides presented by less polymorphic^7^ receptors such as CD1a^8,9^, CD1b^10,11^, CD1c^12,13^, CD1d^14,15^, HLA-E^16^, or MHC-related protein 1 (MR1)^17,18^. Furthermore, some of these cell subsets can be activated in a TCR-independent fashion, *i.e.,* in the absence of their target antigen or in an innate-like manner, using cytokines such as IL-12 and IL-18^19,20^ which cause these cells to release large quantities of cytokines for example INF-γ and TNF-α^21^.

Several subtypes of unconventional T cells have been identified such as mucosal-associated invariant T (MAIT) cells^1,22^, natural killer T cells (NKT)^23–25^, and germline-encoded mycoyl-reactive (GEM) T^11^ cells, with MAIT cells being one of the most well-characterized groups. MAIT cells are characterized by a semi-invariant Vα-7.2^+^ (*TCRAV01-02*) TCR-alpha chain that pairs with a more diverse set of beta-chains, albeit with some preferential V-gene usage such as Vβ13 (*TCRBV06*) family and Vβ2 (*TCRBV20*) family to form a TCR that recognizes bacterial vitamin B2 metabolites presented on MR1 protein^22,26,27^. Although the full antigenic repertoire of MAIT cells has not been determined, some cognate antigens have been identified such as 7-hydroxy-6-methyl-8-D-ribityllumazine (RL-6-Me-7-OH)^28,29^, 5-(2-oxoethylideneamino)-6-D-ribitylaminouracil (5-OE-RU) ^30^ and 5-(2-oxopropylideneamino)-6-D-ribitylaminouracil (5-OP-RU)^30^. In peripheral blood, MAIT abundance ranges from ∼1 – 10% of T cells^31^. As their name implies, MAIT cells also localize to body barriers and tissues, with an abundance of ∼20 – 40% in the liver^31^, 1.2-2.5% in the gut ileum, and 1-10% in the colon^32^.

Given the ability of MAIT cells to bridge innate and adaptive immunity and their dominant presence in different tissues and body compartments, they have been implicated in a wide spectrum of physiological and pathological processes. MR1 deficient mice have been shown to develop a worse disease course in infection models of *K. pneumoniae* and *E. coli* relative to wild-type mice^33–36^. Similarly, MAIT cells have been implicated in other types of bacterial infections such as *L. longbeachae*^35,37,38^, *M. tuberculosis (Mtb)*^39,40^, and *F. tularensis*^40–42^. Viral infection can also trigger MAIT activation, although this is mediated in a TCR-independent manner primarily using IL-18^43,44^. Besides controlling infectious agents, MAIT cells have been implicated in autoimmune and inflammatory diseases, for example, ∼ 5% of CD8^+^ T cells identified in Multiple Sclerosis brain lesions are MAIT cells (defined as Vα7.2^+^CD161^+^ cells)^45,46^. Furthermore, in inflammatory bowel disease, the frequency of MAIT cells in peripheral blood has been repeatedly shown to be decreased relative to healthy individuals^46–48^.

In most, if not all cases, MAIT cells are identified using staining of different cell markers such as CD161, Vα-7.2, or MR1 tetramers loaded with known antigens such as 5-OP-RU. Nonetheless, this has three limitations, first, it limits the starting biological material to intact cells which is not always available or accessible, second, it has a low throughput and cannot be easily scaled to a large number of samples and lastly, it is labor intensive and time consuming. To mitigate these limitations, we developed *sequence-to-MAIT (Seq2MAIT)* which is a deep learning framework that can be used to identify MAIT cells directly in T cell repertoire sequencing datasets. This enables Seq2MAIT to be used with existing datasets and with any sample where DNA or RNA is available. As a result, *Seq2MAIT* can be used to estimate the percentage of MAIT cells in different tissues under different physiological and pathological conditions, such as infection, inflammation, aging as well as different cancers. This would shed some light on the contribution of MAIT cells to these different conditions as well as contributing to the development of novel therapeutic targets. Beyond this, *Seq2MAIT* can also be used to annotate bulk TCR-Seq datasets, as *Seq2MAIT* can assign a probability score of MAIT identity to each clonotype. This can be used to predict the source of many disease-associated clonotypes^49,50^, *i.e.* estimate the likelihood that these disease-associated clonotypes are derived from MAIT cells, which leads to the development of a better understanding of the pathophysiological mechanisms involved in the disease. Here, we describe the development of *Seq2MAIT* and benchmark its performance on multiple independent test datasets.

## Materials and Methods

### I. Sample collection and cohort description

TCR and HLA sequence data from human samples used for these studies were aggregated from several independent study collections described below. All necessary patient/participant consent has been obtained for each study and the appropriate institutional forms have been archived. PBMC used for sorted repertoire experiments were collected either by DLS (Discovery Life Sciences, Huntsville, AL) under Protocol DLS13 for collection of clinical samples or by Bloodworks Northwest (Seattle, WA). Volunteer donors were consented and collected under the Bloodworks Research Donor Collection Protocol BT001. An independent cohort of cells sorted for CD8^+^, CD161^+^, and Vα-7.2^+^ was included for additional validation^51^.

### II. Isolating and sorting MAIT cells using MR1 tetramers

PBMCs from 20 donors were stained with MR1 tetramer and sorted for repertoire sequencing. To enhance tetramer staining PBMCs were treated with the protein kinase inhibitor Dasatinib at 50nM for 10 minutes at 37C. Cells were then treated with Fc receptor blocking solution (Human TruStain FcX, Biolegend) for 5 minutes at RT followed by MR1 tetramer staining (T-Select Human MR1 Tetramer v2-PE, MBL). The tetramer was pre-loaded with 5-OP-RU and stored according to the manufacturer’s protocol. MR1 tetramer staining was performed for 40 minutes at 4C. Midway through the tetramer stain PE-Cy7-TCR Vα7.2 antibody (clone 3C10, Biolegend) was added to the cells for the remainder of the stain. To enrich for stained cells and decrease sort time, cells were washed after that stained with anti-PE ultrapure microbeads (Miltenyi Biotec) for 15 minutes at 4C. After washing a small aliquot was taken for pre-enrichment flow analysis and the remaining cells were loaded onto magnetic columns (MS or LS columns, Miltenyi Biotec). PE and PE-Cy7 stained cells were captured on the column and subsequently eluted after washing away unstained cells. After elution, the cells were stained for 20min at 4C with an antibody cocktail that included CD3-BV786, CD4-BV510, CD8-FITC, CD161-BV421, CD14-PerCP_Cy5.5 and CD19_PerCP_Cy5.5. Cells were washed and resuspended in MACS buffer and BD ViaProbe Cell Viability Solution (BD Biosciences) was added prior to loading onto the BD FACSMelody. Populations sorted: ViaProbe-/CD19-/CD14-/CD3^+^/Vα7.2^+^and-/MR1-5OP-RU^+^ and ViaProbe-/CD19-/CD14-/CD3^+^/Vα7.2^+^/MR1-5OP-RU-. In addition to the sorted samples, there were 19 PBMC donors analyzed by flow cytometry using the same reagents. For flow analysis alone, magnetic bead enrichment was not performed, and all antibodies were added at the same time after the tetramer stain. The MR1 bound fraction of the total CD3^+^ population was calculated using the pre-enrichment sample for sorted donors as well as the samples analyzed by flow for unsorted donors. Calculations were performed using FloJo Software.

### III. Repertoire sequencing

All sorted cells, which ranged from 9.5k to 1.1million, were sent for sequencing. Two million PBMCs from each of the 20 sorted donors were also sequenced along with five million PBMCs from 19 unsorted donors. RNA was isolated from the sorted samples and both RNA and gDNA were isolated from the matched PBMCs. The sorted samples and their matched PBMCs were then divided in two and sent for TCR-α and TCR-β sequencing using the ImmunoSEQ assay at Adaptive Biotechnologies and TCR-β/TCR-α/TCR-δ sequencing was performed in our high throughput R&D lab. The TCR-β sequencing leveraged cDNA for the sorted samples and their matched PBMCs. The TCRβ/TCRα/TCRδ sequencing leveraged cDNA for the sorted samples and gDNA for their matched PBMCs. The 19 PBMCs from unsorted donors leveraged gDNA only for both sequencing assays.

### IV. Model architecture

*Seq2MAIT* is a transformer-based deep learning architecture^52^ that is used to model the probability that a TCR-α or TCR-β chain is derived from a MAIT’s TCR. Each clonotype derived from an alpha or a beta chain is represented using three components, namely, the amino acid sequence of the <u>c</u>omplementary <u>d</u>etermining <u>r</u>egion 3 (CDR3), the V gene, and the J gene that recombined to form this TCR-α or TCR-β chain. *Seq2MAIT* has three input layers the first and the third layer receives tokenized V and J genes, *i.e.* a numerical translation of each gene into an integer, *e.g.*, the gene *TCRBV06-02* is represented as integer 3. However, the second input layer receives the padded and tokenized CDR3 sequence, in which each CDR3 shorter than 20 is padded with zeros, subsequently, each amino acid is translated or mapped to a specific integer, for example, the amino acid lysine (L) is mapped to integer 4. Hence, the model expects TCR chains to be numerically encoded as a tuple of three vectors, the first is the numerically encoded representation of the V gene which is a vector of length 1. The second vector is the numerical representation of the CDR3-sequence, which is a vector of length 20 and lastly, the third vector is a numerical representation of the J-gene, again, a vector of length 1.

After feeding each vector to its corresponding input layer, each token is projected to a learned embedding space of eight dimensions^53^. In the case of V and J genes, this is achieved using an embedding layer while the CDR3 is encoded using a positional encoding embedding layer. It is also worth mentioning that each of the three inputs has its own embedding, for example, there is a dedicated embedding layer for V genes that is not used to embed the CDR3 or the J gene and vice versa. After embedding, the TCR chain is presented as a tuple of three matrices, the first is the v gene (1 × 8), the second is the CDR3 region (20 × 8) and the third is the J gene (1 × 8). These three matrices are then concatenated to generate a 22 × 8 matrix that contains a learned representation of the provided TCR chain. The matrix representation is then forwarded to a transformer-encoded layer to learn dependencies and interactions between the V, J, and CDR3 tokens. Subsequently, the output of the transformer layer is reduced using a max-pooling layer and is then forwarded to a fully connected layer that further processes the generated representation by the transformer layer and then produces a value between zero and one that represents the probability that the input TCR chain belongs to a MAIT’s TCR.

### V. Training and implementation

*Seq2MAIT* was implemented using Keras^54^ and TensorFlow^55^ and the prediction problem was formulated as a classification task in which the model learns features and representations to classify an input sequence (TCR-α or TCR-β) into MAIT (positive class) or non-MAIT (negative class). To do this, the set of unique TCR sequences identified from MR1 sorted cells was defined as the positive class, and an equal number of sequences were randomly selected from unsorted repertoires (bulk TCR-Seq) and labeled as the negative class. In case, a TCR sequence was present in the set of positive and negative examples, it was removed from the set of negative examples and a replacement sequence from the unsorted repertoire pool was added. Thus, an equal number of MAIT (MR1 binding T cells) and non-MAIT (randomly selected from unsorted repertoire without overlapping with the MAIT set) TCRs were selected. Subsequently, labels were created for each group, with positives represented as ones and negatives represented as zeros. Lastly, the examples were shuffled and numerically encoded as described above. *Seq2MAIT* was trained using Adam^56^ as an optimizer with the default learning rate in batches of size 4,096 examples for 100 epochs. Binary cross entropy was used as a loss function to optimize *Seq2MAIT’s* weights with equal weight for positive and negative classes.

## Results

### Study design

We started our analysis by obtaining blood samples from a cohort of 20 healthy individuals (**Fig. 1A**), subsequently, inducing with MR1-5-OP-RU tetramers was used to isolate MAIT cells (defined as CD3^+^TCRVα7.2^+^MR1^+^) (**Materials and Methods**) from the donors’ PBMCs (**Fig. 1B-1C**). RNA and DNA were extracted from the isolated MAIT cells and the unsorted repertoire (**Fig. 1D**). Subsequently, immunoSEQ was conducted to characterize the TCR-Seq repertoire of the MAIT cells and of total T cells in the blood (**Materials and Methods).** The sorted MAIT TCR repertoire was analyzed and computationally compared to the unsorted population of T cells (**Fig. 1E**). Next, a training dataset was constructed which contained the MAIT’s TCR-α/β repertoires and an equivalent number of clonotypes derived from the unsorted T cell repertoire (**Materials and Methods**). Model performance was measured and evaluated on independent test datasets that were not used in either the training of the models or in optimizing their architecture (**Fig. 1F**).

**Figure 1:**
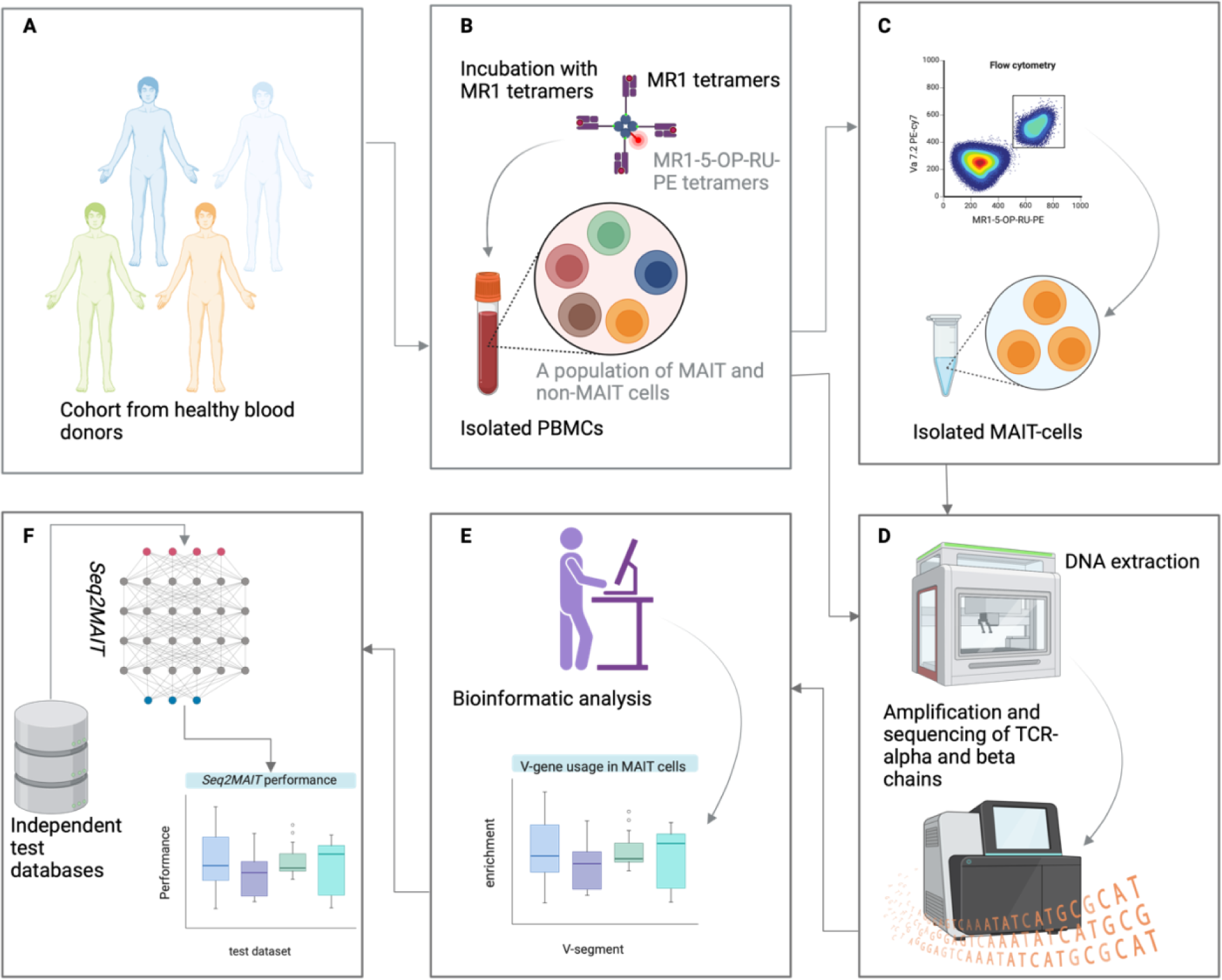
A schematic representation of the study design. **(A)** Assembly of a cohort of healthy donors to isolate blood samples. **(B)** Incubation of isolated samples with MR1 tetramers and other markers such as anti-alpha 7.2 antibodies to isolate this population of cells. **(C)** Isolation of MAIT cells using FACS. **(D)** DNA was extracted from the unsorted population of cells as well as the sorted MR1^+^ cells, subsequently, TCR-α and TCR-β were sequenced using immunoSEQ. **(E)** Bioinformatic and computational analysis of the MAIT cells, *i.e.* MR1^+^ sorted cells, gene usage relative to unsorted repertoire. **(F)** Development and training of Seq2MAIT using the dataset curated and assembled in **(E)** as well as the testing of the model on different independent test datasets.

### Characterizing the alpha and the beta-chains of MR1 sorted cells

We started our analysis by looking at the preferential usage of different V and J genes, in the alpha and beta chain of MAIT cells relative to unsorted repertoire. The unsorted T cell repertoires were used to estimate the background level of gene usage in peripheral blood. Unsurprisingly, the alpha-chain of MR1-sorted cells showed a strong preference for using the *TCRAV01-02* gene, *i.e.* α-7.2, which was used in >30% of all MAIT cells (**Fig. 2A**). This confirms what has been identified previously about the biased usage of *TCRAV01-02* gene in MAIT cells^57^. A weaker preference toward specific J genes (*e.g. TCRJ33-01* and *TCRJ20-01* (**Fig. 2C**)) could also be observed, which agrees with what has been previously shown^57,58^. Interestingly, the degree of J gene preference was much weaker than that observed at the V-gene level with the most used J gene, *TCRJ33-01*, being used by less than 10% of all clonotypes identified from MR1 sorted cells.

**Figure 2:**
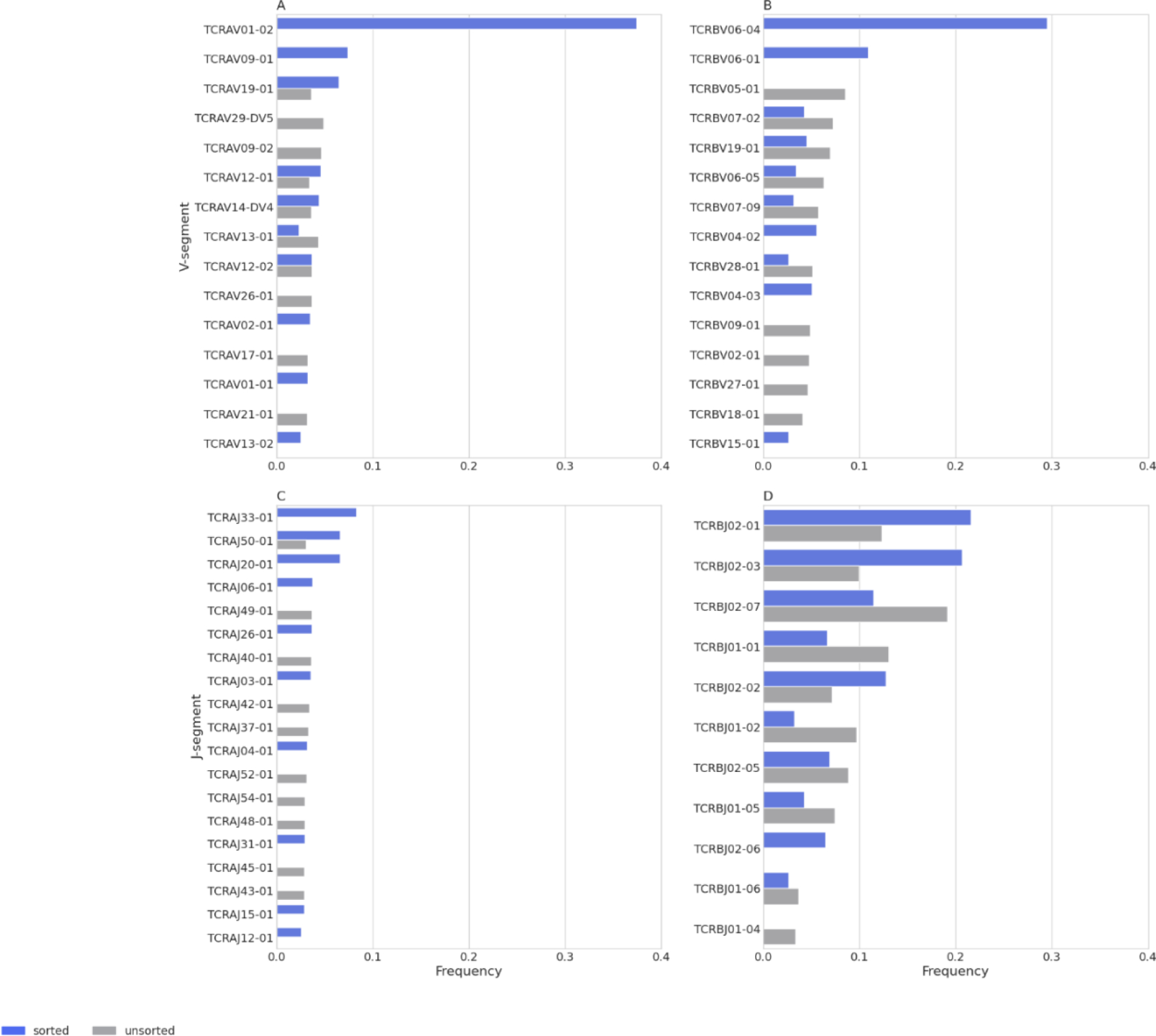
Differential usage of V and J genes in MR1-sorted T cells (MAIT) relative to unsorted repertoires. Plots are based on 9,816 MR1-sorted TCR-α sequences, 709,734 unsorted TCR-α sequences, 14,284 MR1-sorted TCR-β sequences, and 931,612 unsorted TCR-β sequences. The plots show the 10 most commonly found genes in the category of **(A)** TCR-α V gene usage **(B)** TCR-β V gene usage **(C)** TCR-α J gene usage **(D)** TCR-β J gene usage.

At the TCR-β chain, MAIT cells have been shown to exhibit a strong preference for using genes of the TCRBV06 family, such as *TCRBV06-04* and *TCRBV06-01* as well as the TCRBV04 family (*e.g.*, *TCRBV04-01* and *TCRBV04-03*)^57,58^. This was strongly seen in our dataset where ∼30% and ∼10% of all MAIT’s clonotypes were derived from *TCRBV06-04* and *TCRBV06-01 V* genes, respectively (**Fig. 2B**). These usages also agree with study from Garner and colleagues^58^ which showed a preferential usage of *TCRBV06-01 and TCRBV06-04* in MAIT cells using single-cell RNA sequencing of MAIT cells (defined as: CD8^+^CCR7^−^MR1-5-OP-RU^+^). A weaker preference can be observed at the J gene level, with *TCRBJ02-01* and *TCRBJ02-03* being the most utilized genes (**Fig.2D**).

MAIT cells are expected to be a group of cells targeting vitamin B metabolites presented by MR1 protein, thereby sharing similar and public TCR sequences. To quantify the degree of publicity of MAIT TCRs we looked at the fraction of MAIT TCRs that is shared among an increasing number of individuals (**Fig. S2A**) and investigated the relationship between the degree of publicity and the generation probability, *i.e.* the likelihood of generating a specific T cell clonotype that is a specific V(D)J rearrangement^59^, estimated using OLGA^60^ (**Fig. S2B**). As seen in **Fig. S2A,** the majority of MAIT TCR sequences are private, *i.e.*, are observed only in a single donor. Furthermore, there was an inverse relationship between the generation probability and the degree of publicity. We also sampled an equal number of clonotypes from the unsorted repertoires and compared their generation probability to that of MAIT cells (MR1 sorted cells). As seen in **Fig. S2C**, although the majority of MAIT cells are private with a wide range of generation probabilities, they tend to have a higher generation probability relative to the unsorted repertoire which contains a large fraction of conventional α/β T cells.

### Seq2MAIT can accurately distinguish MAIT from non-MAIT cells

Given the observed diversity in the TCR sequences of MAIT cells and the lack of a clear motif that distinguishes these cells from other T cell subsets, we sought to develop a deep-learning model to address this problem. The developed model will extract generalizable subtle features from the TCR sequences of MR1-sorted cells, enabling the model to discriminate between MR1-sorted cells (MAIT cells) and other T cell populations based on either TCR-α or TCR-β sequences.

To this end, we developed the sequence-to-MAIT model (*Seq2MAIT*) which is a transformer-based model that receives TCR-α or TCR-β chain clonotypes as an input and returns the probability that the provided sequence is derived from a MAIT cell. Most of the observed variation in TCRs is in complementary determining regions (CDRs)^61^ with each of the TCR chains exhibiting 3 CDRs, namely, CDR1, CDR2, and CDR3 (**Fig. 3A**). Studies investigating the interaction between TCRs, and peptide-HLA complexes have shown that CDR1 and CDR2 are involved in interacting with HLA proteins while CDR3 are mainly involved in interacting with the presented peptide^62,63^. CDR1 and CDR2 are germline-encoded in the V gene while the CDR3 is formed due to somatic recombination between the V and the J gene in the case of the TCR-α chain and the V, the J and the D gene in the case of the TCR-β chain in addition to random nucleotide insertions and deletions during the recombination event (**Fig. 3A**). Hence, we represented a TCR chain as a unique combination of a V-gene, a J-gene, and the amino acid sequence of the CDR3 (**Fig. 3A**). After selecting this representation, we assigned each V and J gene a unique integer that represents its identity numerically, *i.e.* a V or J token, and encoding each amino acid in the CDR3 as a unique integer representing the code of the amino acid, *i.e.* amino acid token (**Materials and Methods).**

**Figure 3:**
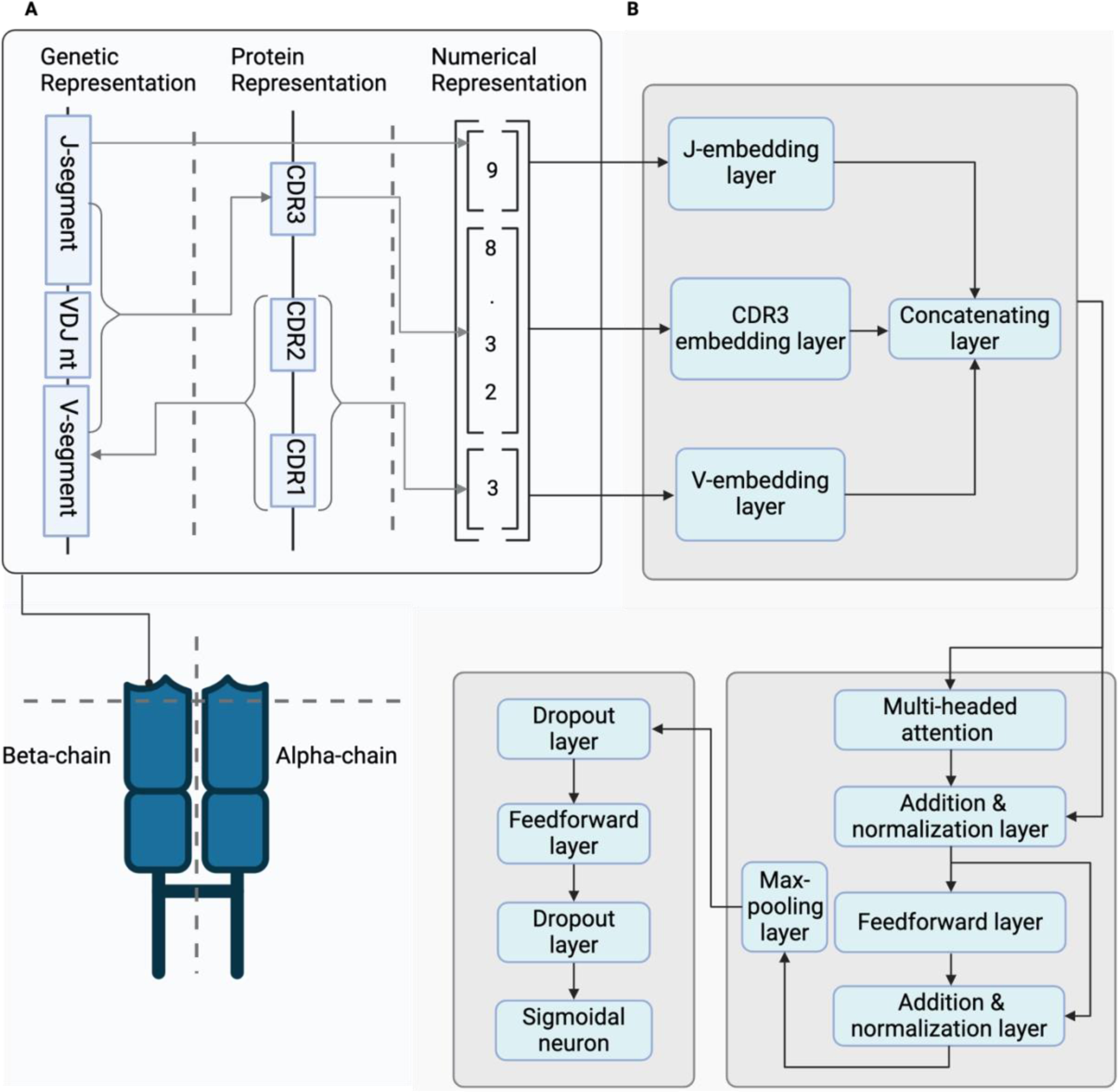
The numerical representation of a TCR and the layout of Seq2MAIT. **(A)** The representation and numerical encoding of TCR chains. **(B)** The layout of Seq2MAIT is built using three modular parts or components: (i) an embedding part responsible for generating a learned representation of the encoded TCR representation, (ii) a self-attention component responsible for learning interactions and dependencies between the V and J genes and the CDR3. Lastly, a decision-making component is responsible for calculating the probability that a sequence is a MAIT sequence given its learned and processed representation.

The architecture of *Seq2MAIT* was built to support this encoded representation of TCR chains where the model receives three inputs, representing the code of each V and J gene as well as the code of each CDR3 from either TCR-α or TCR-β chains. A learned embedding layer is used to represent, in a task-specific manner, the corresponding V and J gene as well as the CDR3 (**Fig. 3B**), followed by a transformer encoder layer^52^ to learn dependencies and interactions between these components of the TCR chains (**Fig. 3B**). This generates an information-rich representation of the chain that is forwarded to feedforward layers, *i.e.* fully connected neural networks, and after that to a sigmoidal neuron to calculate the probability that a given sequence is derived from a MAIT cell (**Fig. 3B; Materials and Methods**).

Given the observed diversity at the TCR-α and the TCR-β chains of MAIT cells, we trained two versions of *Seq2MAIT*, each corresponding to one of the chains, *i.e.* training a model on the TCR-α chain and a model on the TCR-β chain separately. As described in the **Materials and Methods,** *Seq2MAIT* was trained on an equal number of negative cases (*i.e.* unsorted repertoire) and positive cases (*i.e.* MR1 sorted clonotypes). To control for the strong bias in the V gene of the alpha-chain of MAIT cells, where most sequences are derived from *TCRAV01-02*, we developed a training dataset composite of MAIT and non-MAIT sequences derived from *TCRAV01-02* clonotypes. This enabled us to build a classifier that can predict whether a *TCRAV01-02* clonotype is derived from a MAIT cell or not without any bias introduced by the difference in the *TCRAV01-02* gene usage in MAIT relative to unsorted T cells.

The TCR-α model was trained on 3,671 MAIT sequences with a matching number of controls for each chain, we tested the generality of the model on a test dataset composite of 366 MR1^+^ TCRs, *i.e.* MAIT, and 369 non-MR1 TCRs sampled from unsorted repertoires. Seq2MAIT had an <u>a</u>rea <u>u</u>nder the receiver <u>o</u>perator <u>c</u>urve, AU[ROC] of ∼ 0.76 (**Fig. 4A**) and an <u>a</u>rea <u>u</u>nder the <u>p</u>recision-recall curve AU[PR] of ∼0.79 (**Fig. 4B**). Regarding TCR-β sequences, we trained a *Seq2MAIT* model on 10,975 MR1-sorted clonotypes (∼81% of the full dataset) and equivalent matched controls. We optimized its hyperparameters using a validation dataset composite of 1,248 MR1-sorted clonotypes (∼9% of the full dataset) and equivalent matched controls. Lastly, its performance was measured on the test dataset which contains 1,381 MR1-sorted clonotypes (∼10% of the full dataset) and equivalent matched controls. As seen in **Fig. 4C and 4D**, *Seq2MAIT* can discriminate between MAIT-derived and non-MAIT TCR-β chains with an AU[ROC] of ∼0.875 and AU[PR] of ∼0.882.

**Figure 4:**
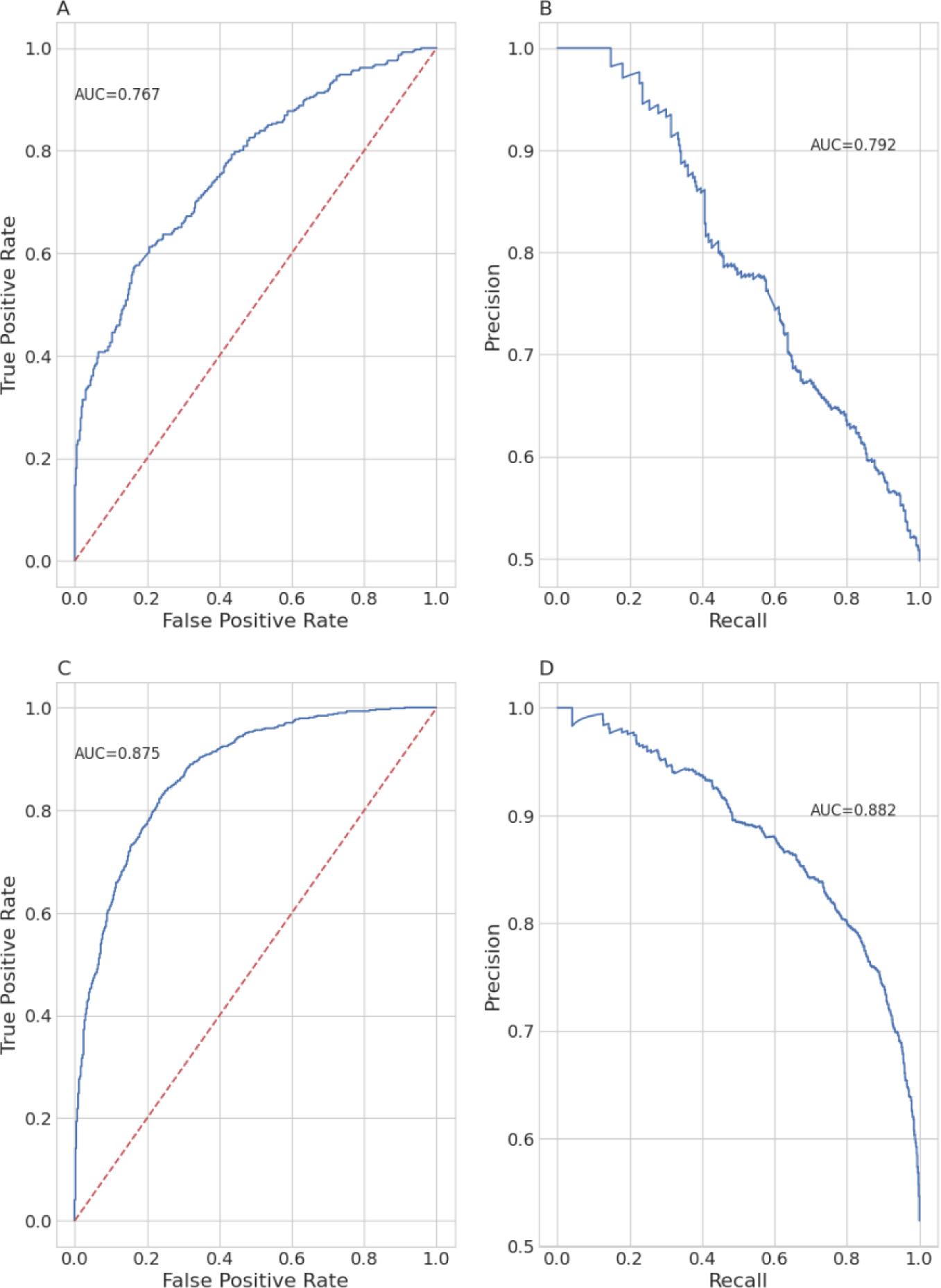
The performance of the TCR-α and TCR-β models on unseen test datasets. **(A)** represents the receiver operator curve and **(B)** represents the precision-recall curve for the predictions of the TCR-α model on 366 MR1 TCRs and 369 non-MR1 binding, *i.e.* unsorted, TCRs both of which are derived from *TCRVA01-02* gene. **(C)** is the receiver operator curve and **(D)** is the precision-recall curve for the predictions of the *TCR-β* model on 1,381 MR1 TCRs and 1,340 non-MR1 TCRs.

To characterize the impact of different parts of the input on the overall performance, we masked different combinations of the inputs, for both TCR-α and TCR-β chains, and quantified the impact on the model performance (**Fig. S3**). As seen in the figure, the models achieve their highest performance accuracy when conditioning the predictions on encoded V and J genes as well as CDR3s, while zero-masking all inputs, produces a random model. Beyond these two trivial cases, we observed a substantial degree of information-leaking between these three components, for example, conditioning the predictions on the V and J gene only or the CDR3 sequences only, did not result in a substantial deterioration of the predictive performance. Given the encouraging performance of the *Seq2MAIT* TCR-β model and the widespread availability of TCR-β datasets, we focused our efforts on further optimizing the *Seq2MAIT* model for identifying MAIT’s TCR-β sequences.

To investigate the generalizability of *Seq2MAIT* on datasets generated under different experimental conditions we utilized the dataset of MAIT cells (defined as CD8^+^CD161^+^Vα-7.2^+^) recently published by Williams and colleagues^51^. Taking only the MAIT sorted sequences from this dataset, we started by looking at the distribution of the model’s scores for these clonotypes. As seen in **Fig. 5A**, for most TCR-β chains (∼84% of all TCR-β chains in the dataset) the estimated MAIT probabilities (*i.e. Seq2MAIT* scores) were above 0.5 indicating the generalizable performance of the model. To gain a better understanding of cases where *Seq2MAIT* predicted a low score for MAIT TCR-β chains, we looked at the score distribution within clonotypes derived from different V genes. As seen in **Fig. 5B**, the model scores varied largely between sequences bearing different V genes. Members of the TCRBV06 family, such as *TCRBV06-01* and *TCRBV06-04* which are preferentially used by MAIT cells, are estimated to have a higher likelihood of being MAIT sequences, while families such as TCRBV02 generally have a low score*, i.e.* a low-probability of being MAIT. Given that V genes with low scores are generally not reported in the literature to be preferentially used by MAIT cells, our findings might reflect noise in this test dataset introduced by using a different definition of MAIT cells (CD8^+^CD161^+^Vα-7.2^+^,) which contains MR1-bindings and non-MR1 binding T cells.

**Figure 5:**
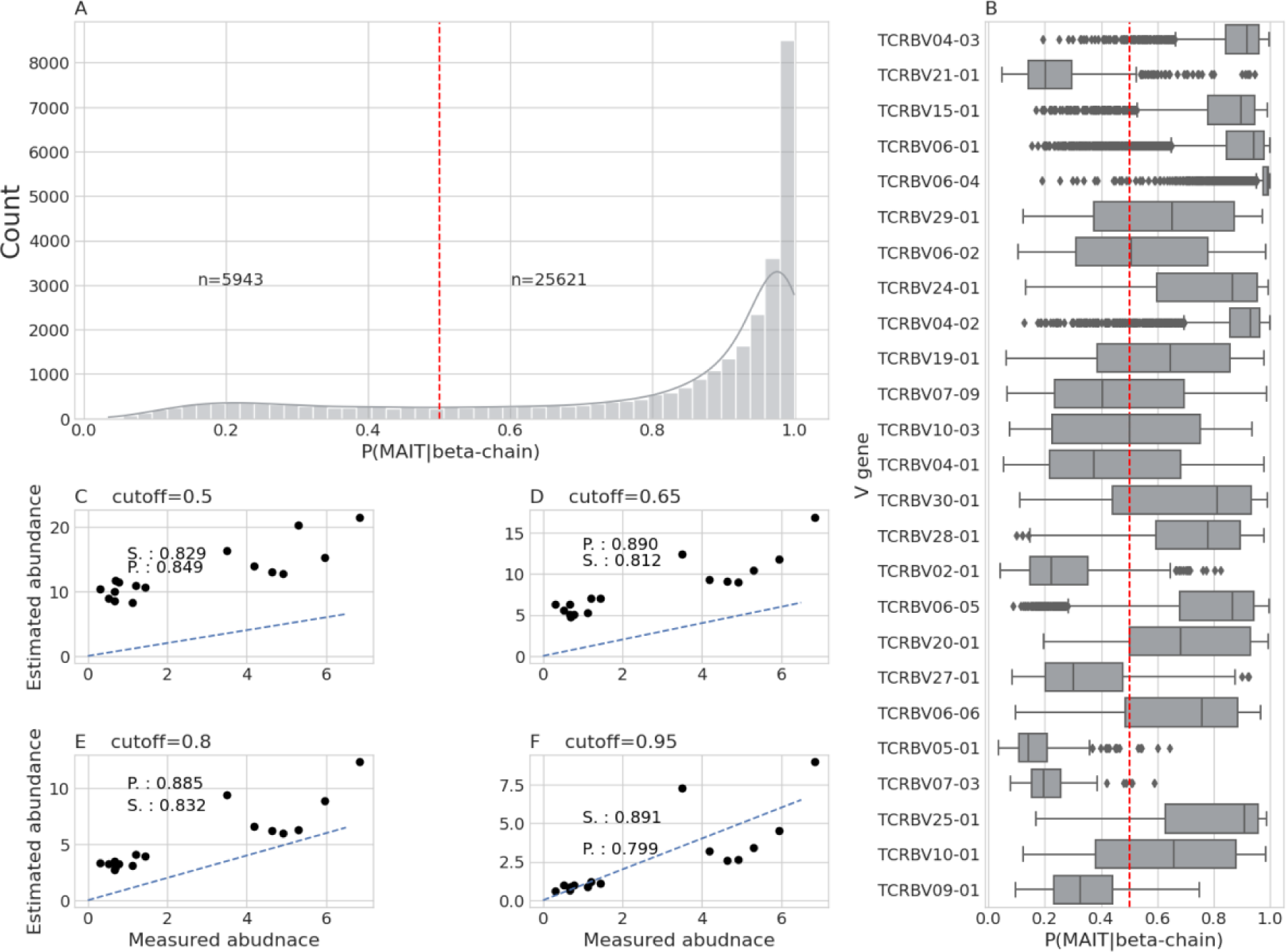
Model performance on independent test datasets. **(A)** Distribution of models’ scores on a publicly available test dataset of MAIT cells. **(B)** Distribution of the model’s scores across the TCR-β chain belonging to different V genes. **(C)** Correlation between the estimated MAIT abundance using *Seq2MAIT* and the experimentally measured abundance using a cutoff of 0.5, while **(D)** shows the correlation using a cutoff of 0.65, **(E)** shows 0.8 as a cutoff value, and lastly **(F)** shows the correlation between the estimated abundance and the experimentally measured abundance using 0.95 as a cutoff value.

Lastly, we were interested in assessing the ability of *Seq2MAIT* to quantify the abundance of MAIT cells in the blood. To this end, we generated a dataset in which the TCR-β of unsorted T cells from blood were sequenced, and the abundance of MAIT cells was quantified using flow cytometry on MR1-tetramers stains (**Materials and Methods**). To quantify the abundance of MAIT cells computationally, we first utilized *Seq2MAIT* to identify MAIT clonotypes from the unsorted repertoire data using different cutoff values (**Fig. 5C-5F**). Subsequently, MAIT cell abundance was estimated by summing the template count of each MAIT clonotype and dividing this sum by the total template count of the unsorted repertoire. As seen in the figure, there is a strong positive correlation between the computationally estimated abundance of MAIT cells using *Seq2MAIT* and the experimentally measured values. With lower cutoffs (*i.e.* only counting sequences with larger predicted scores), the estimated abundance is generally much higher than experimentally measured values **(Fig. 5C),** however, as the cutoff increases, the predicted MAIT abundance approaches experimentally measured values **(Fig. 5E-5F)**.

## Discussion

MAIT cells are a group of unconventional T cells that are characterized by a semi-invariant TCR-α chain and can recognize vitamin B metabolites presented on MR1 proteins^22,26,27^. Despite their abundance and relevance to different physiological and pathological processes, characterizing MAIT cells has proven to be a difficult task because MAIT cells are primarily defined by flow cytometry, either using MR1 tetramer staining or using a combination of other markers such as CD161 and Vα7.2. Although flow cytometry can provide valuable insights about the abundance of MAIT cells and their phenotype, it is time and labor-intensive, requires access to viable cells, and has a low to medium throughput.

Here, we tried to address this problem by developing *Seq2MAIT* which is a deep-learning model for identifying MAIT cells in bulk TCR repertoire sequencing datasets. As shown above, *Seq2MAIT* can accurately identify the TCR-α and TCR-β chains of MAIT cells with an accuracy of around 80% across different datasets. This enables *Seq2MAIT* to be applied to the hundreds of thousands of repertoires generated over the last two decades. *Seq2MAIT* also enables the abundance of MAIT cells to be quantified in any biomaterial where DNA is available, effectively bypassing the need for living cells and enabling MAIT cell identification in, virtually, any sample.

Despite all the advantages of *Seq2MAIT* it still has several limitations, first, it only enables the identification of MAIT cells without any information regarding other phenotypic markers. For example, with live-cell-based methods, cells can be stained with different markers to identify their phenotypic properties such as activation markers and cytokine receptors, *e.g.* IL-17. This becomes apparent when identifying the subset of MAIT cells, *e.g.* MAIT 1 vs MAIT 17^58,64^, is critical for understanding the biological process under investigation. Hence, *Seq2MAIT* is not intended as a replacement for flow-cytometry-based methods but as a complementary method that can be used to identify MAIT cells in samples where living cells are not available or to prioritize samples for further characterization using flow-cytometry or scRNA-Seq based methods.

*Seq2MAIT* can be categorized as a specialized case of TCR-antigenicity prediction models, such as NetTCR^65^, NetTCR-2.0^66^, and *DeepAIR*^67^, which aim at predicting the interaction between a specific TCR and a given peptide-HLA complex. This is a much harder problem because of the high allelic diversity of HLA proteins relative to the monomorphic MR1 protein as well as due to the large antigenic space of antigenic peptides that are presented on HLA proteins relative to the few vitamin B metabolites presented on MR1 proteins. Nonetheless, the same experimental and modeling strategies used here can be utilized for other non-conventional T cells such as NKT cells, GEM cells, CD1a-restricted cells, *et cetera*. We envision that another version of *Seq2MAIT* will be developed in the foreseeable future to identify and characterize all unconventional T cells in bulk TCR-Seq data. Given that these cells represent a substantial fraction of the T cell population in different tissues and organs such as the liver, gut, and respiratory system, studying their abundance and response to different stimuli would contribute to a better understanding of their contribution to different physiological and pathological processes.

Although, to the best of our knowledge, *Seq2MAIT* is the first deep learning framework that can be used to accurately identify MAIT cells from sequencing data, different future extensions can be developed to improve its performance further. Currently, *Seq2MAIT* predicts the probability of MAIT cells solely from a single chain of the TCR (*i.e.* TCR-α or TCR-β), however, extending *Seq2MAIT* to model the paired TCR-α/β chains of MAIT cells, *i.e.* receptor-level instead of chain-level modeling, would enhance our ability to identify MAIT cells. Nonetheless, most repertoire sequencing experiments are still performed at the single chain level, mainly the TCR-β chain, and only a limited number of experimental methods can provide paired TCR-α/β chains such as pairSEQ^68^.

A second future direction to improve the model performance further is to extend and refine the definition of MAIT cells. As discussed above, MAIT cells have different definitions such as CD8^+^CCR7^−^MR1/5-OP-RU^+ 58^ or as CD3^+^CD4^−^TCRγδ^−^CD161^high^TCRAV01-02^+ 69^. These definitions generally capture only a subset of the MAIT cell population, for example, the former definition by Garner *et al.*^58^ only captures CD8^+^ MAIT cells and excludes CD4^+^ MAIT cells. Interestingly, Xiong and colleagues^70^ have recently shown an important role for CD4^+^ MAIT cells in responding to *M. tuberculosis* infection. Furthermore, other subsets of unconventional T cells use α-7.2 as an alpha-chain, for example, germline-encoded mycolyl-reactive (GEM) T cells which are a subset of CD1b restricted cells that recognize the *M. tuberculosis-*derived glycolipid *glucosemonomycolate*^11^. Although MR1 tetramers-based sorting enables the identification of MR1 restricted T cells, *i.e.* MAIT cells, it is only based on a handful of antigens, mainly 5-(2-oxopropylideneamino)-6-d-ribitylaminouracil (5-OP-RU). Thus, MR1-tetramers do not capture the full repertoire of MR1-restricted cells but only sample cells that bind reactively to the antigen loaded on the MR1 protein. As a result, performing the sorting with tetramers loaded with different antigens might enable a more inclusive definition of MAIT cells which would enable a better characterization of these cells and their contribution to different biological processes.

In conclusion, our developed model *Seq2MAIT* has shown a robust ability to identify MAIT cells across different datasets. Future efforts shall focus on extending *Seq2MAIT* to other unconventional T cell populations such as GEM and NKT cells which would enable a better understanding of the role of these cells in different diseases as well as advance our understanding of different immunological processes they are involved in.

## Acknowledgments

The authors would like to thank Jabran Zahid (Microsoft Research Lab, Redmond Campus, WA, USA), Erin Calfee (Adaptive Biotechnologies, Seattle, WA, USA), and James Lord (Benaroya Research Institute, Seattle, WA, USA) for their feedback. We thank the DFG Research Unit 5042 miTarget for supporting H.E. with a travel stipend. Lastly, Figure 1 and Figure 3 were Created with BioRender.com.

## Author contributions

M.P and R.T designed and conceived the study. H.E, M.P and SW analyzed the data and benchmarked the performance of *Seq2MAIT*. H.E developed, implemented, and trained Seq2MAIT. R.B, B.R, J.S, R.T performed the cell-sorting and MAIT cells isolation. M.P, A.F, R.T supervised the study. H.E wrote the manuscript with input from all authors. all authors have edited and approved the final manuscript.

## Supplementary figures

**Figure S1:**
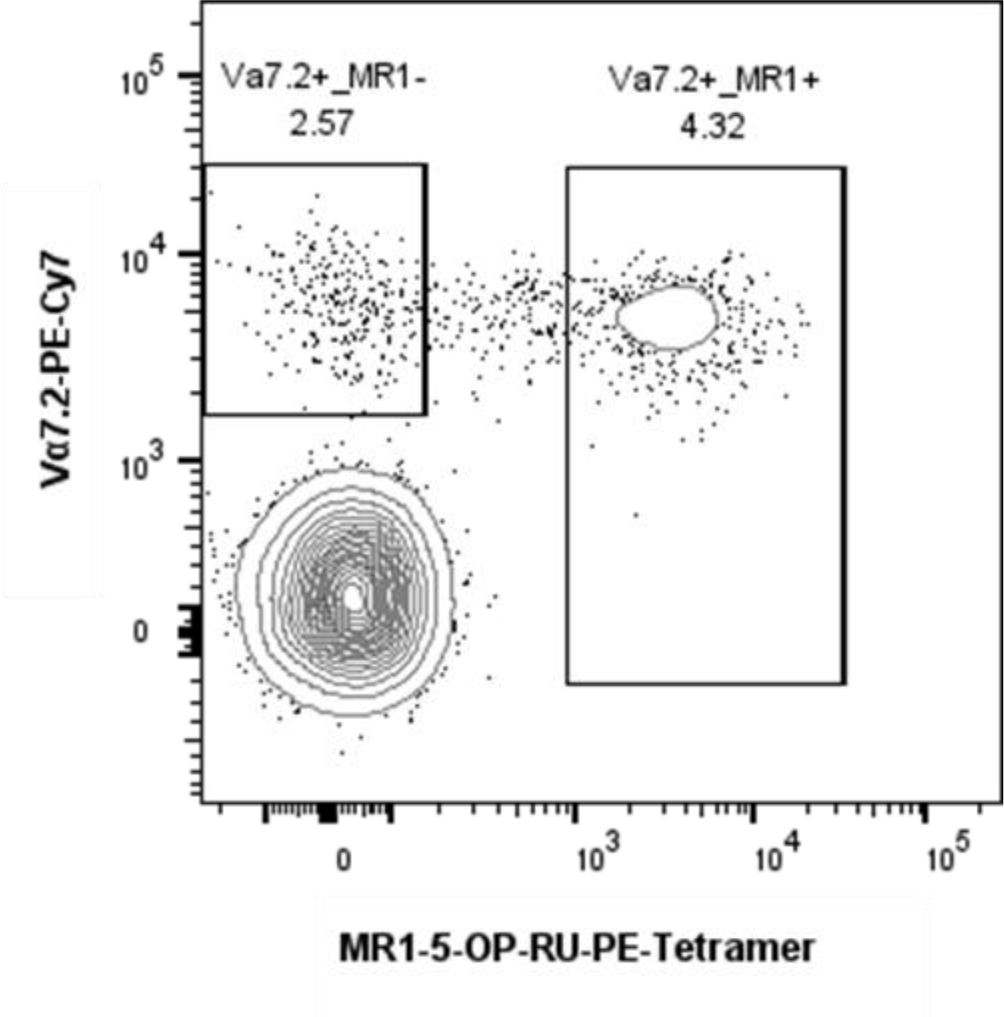
Representative plot of a pre-enrichment sample. Post-enrichment samples were used for sorting, while pre-enrichment samples were used to calculate MAIT frequencies. The gating for the sorts excluded outliers to obtain pure populations. Sorted cells in this plot were first gated on the lymphocyte population in the FSC vs SSC plot then gated to exclude doublets followed by gating on live CD3+ cells.

**Figure S2:**
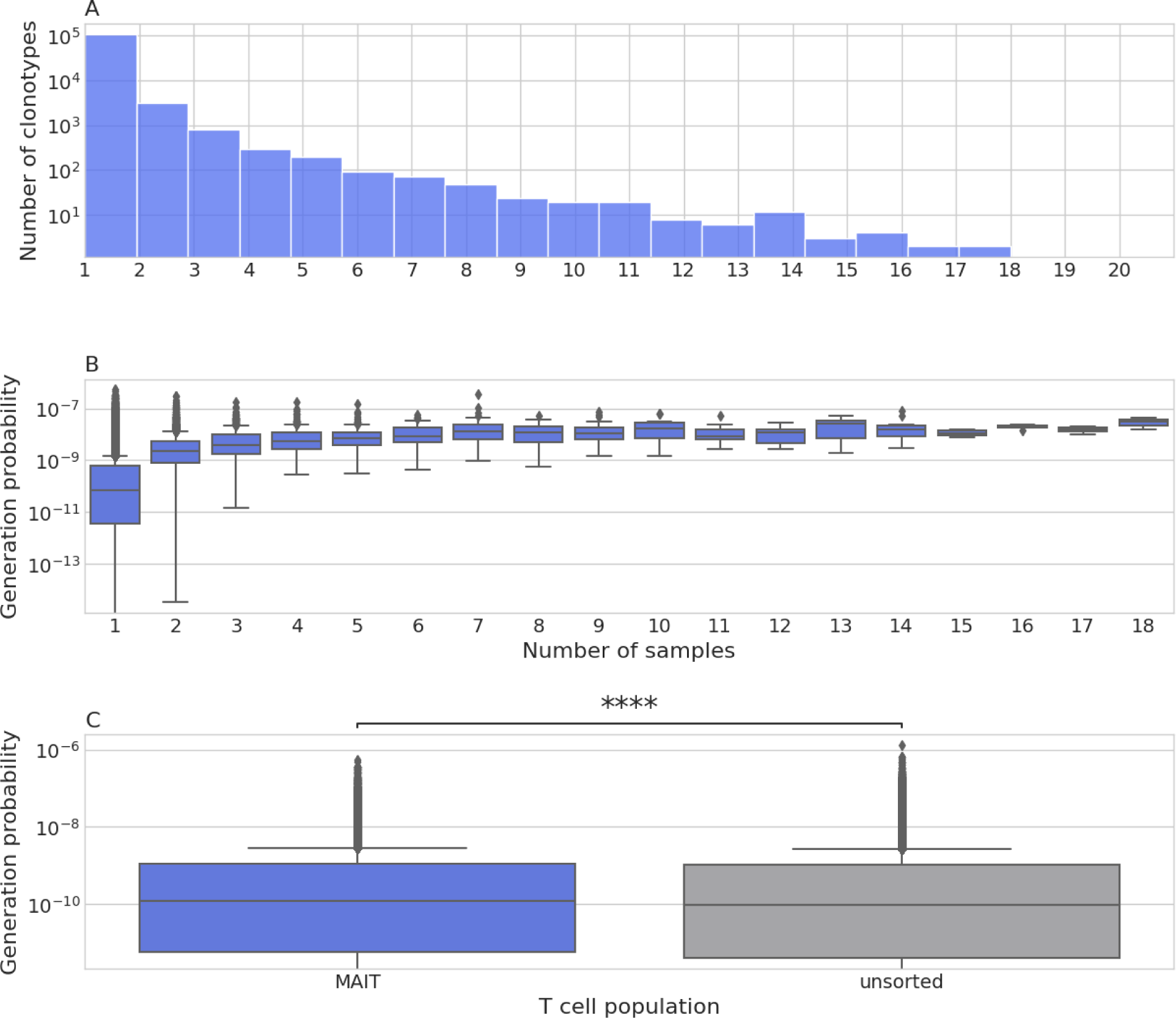
General features of TCR-β chains derived from MR1 sorted cells relative to unsorted repertoires. **(A)** Quantification of the degree of publicity in the TCR-β chain of MR1 sorted cells. The x-axis represents the number of donors, and the y-axis represents the number of MR1-binding TCR-β clonotypes observed in the corresponding number of donors, for example, the first bar represents the number of TCR-β clonotypes observed in only one donor, i.e. private clonotypes. **(B)** Relationship between the number of donors with a specific MR1-binding TCR-β chain and the generation probability of this chain. **(C)** The difference in the generation probability between MR1-binding TCR-β chains relative to the unsorted repertoire.

**Figure S3:**
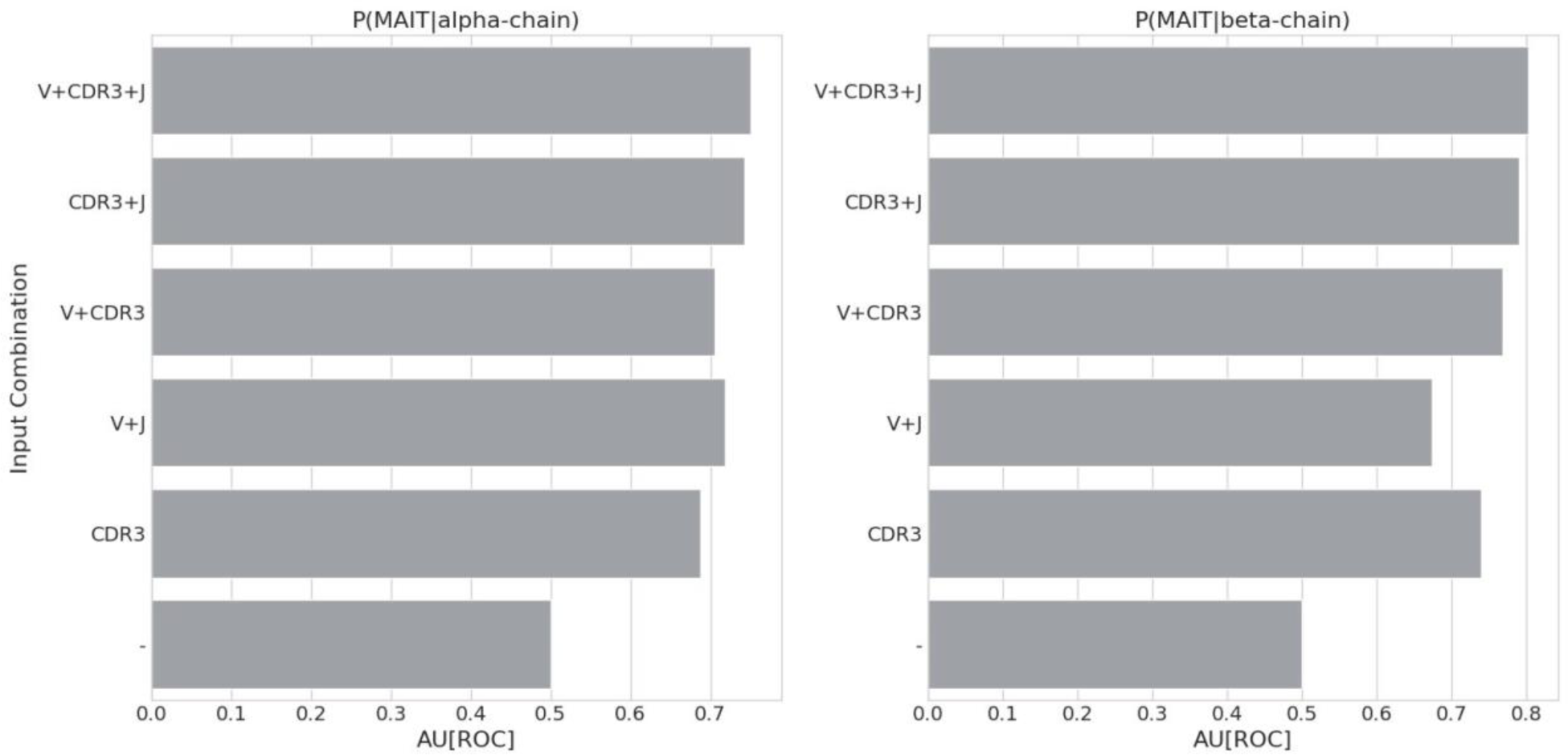
Feature importance analysis where the importance of different inputs on the accuracy of Seq2MAIT is quantified. V and J refer to the V and J genes, respectively, while the CDR3 is the amino acid sequence of the Complementarity-determining region 3.

